# The late ISC pathway interactome reveals mitosomal-cytoplasmic crosstalk in *Giardia intestinalis*

**DOI:** 10.1101/2022.08.01.502261

**Authors:** Alžběta Motyčková, Luboš Voleman, Vladimíra Najdrová, Lenka Marková, Martin Benda, Vít Dohnálek, Natalia Janowicz, Ronald Malych, Róbert Šuťák, Thijs J. G. Ettema, Staffan Svärd, Courtney W. Stairs, Pavel Doležal

## Abstract

Mitochondrial metabolism is entirely dependent on the biosynthesis of the [4Fe-4S] clusters, which are part of the subunits of the respiratory chain. The mitochondrial late ISC pathway mediates the formation of these clusters from simpler [2Fe-2S] molecules and transfers them to client proteins. Here, we characterized the late ISC pathway in one of the simplest mitochondria, mitosomes, of the anaerobic protist *Giardia intestinalis* that lost the respiratory chain and other hallmarks of mitochondria. Identification of the late ISC interactome revealed unexpected involvement of the aerobic marker protein BolA and specific interaction of IscA with the outer mitosomal membrane. Although we confirmed that the synthesis of the Fe-S cluster remained the only metabolic role of mitosomes, we also showed that mitosomes lack client proteins that require the [4Fe-4S] cluster. Instead, by knocking out the *bolA* gene from the *G. intestinalis* genome, we showed that, unlike aerobic mitochondria, the late ISC mitosomal pathway is involved in the assembly of cytosolic [4Fe-4S] clusters. Thus, this work reveals an unexpected link between the formation of mitochondrial and cytosolic [4Fe- 4S] clusters. This may either be a consequence of mitochondrial adaptation to life without oxygen, or it represents a general metabolic coupling that has not been previously observed in the complex mitochondrial metabolism of aerobes.

## INTRODUCTION

*Giardia intestinalis* is a microaerophilic parasitic protist that lives in the epithelium of the small intestine of mammals, where it causes giardiasis (1). It belongs to the Metamonada supergroup of eukaryotes that typically contain mitochondria-related organelles (MRO) that lack organellar genomes and cristae and that are adapted to life with little or no oxygen (2). The so-called mitosomes of *G. intestinalis* are one of the simplest MROs known among eukaryotes, as they contain only a single metabolic pathway for iron-sulfur (Fe-S) cluster synthesis (ISC) (3–5).

Fe-S clusters function as cofactors of proteins (Fe-S proteins) in all living organism. In eukaryotes, they participate in essential biological processes in various compartments such as DNA maintenance in the nucleus, electron transport chains in mitochondria, and protein translation in the cytoplasm (6–8). In humans, about 70 different Fe-S proteins have been identified (7).

In aerobic eukaryotes, the formation of Fe-S clusters for all cellular proteins begins in mitochondria via the activity of the ISC pathway, which can be functionally divided into the early or late acting complex of proteins (9). In ‘classical’ mitochondria (Fig. 1A), the early ISC pathway produces [2Fe-2S] clusters on the scaffold protein IscU (10) via the activity of a complex consisting of cysteine desulfurase IscS (11), its accessory subunit Isd11 (12–14) and an acyl carrier protein (15–17). The actual transfer of sulfur to IscU is facilitated by frataxin (18) and the electrons for cluster formation are provided by reduced ferredoxin (Fdx), which itself is a [2Fe-2S] protein (19). However, the source of iron and the mechanism of iron transfer to the cluster remain elusive. Upon the formation of [2Fe-2S] cluster on IscU, a chaperone complex consisting of Hsp70 and HscB transfers the cluster to glutaredoxin 5 (Grx5) apoprotein (20).

**Figure 1.**
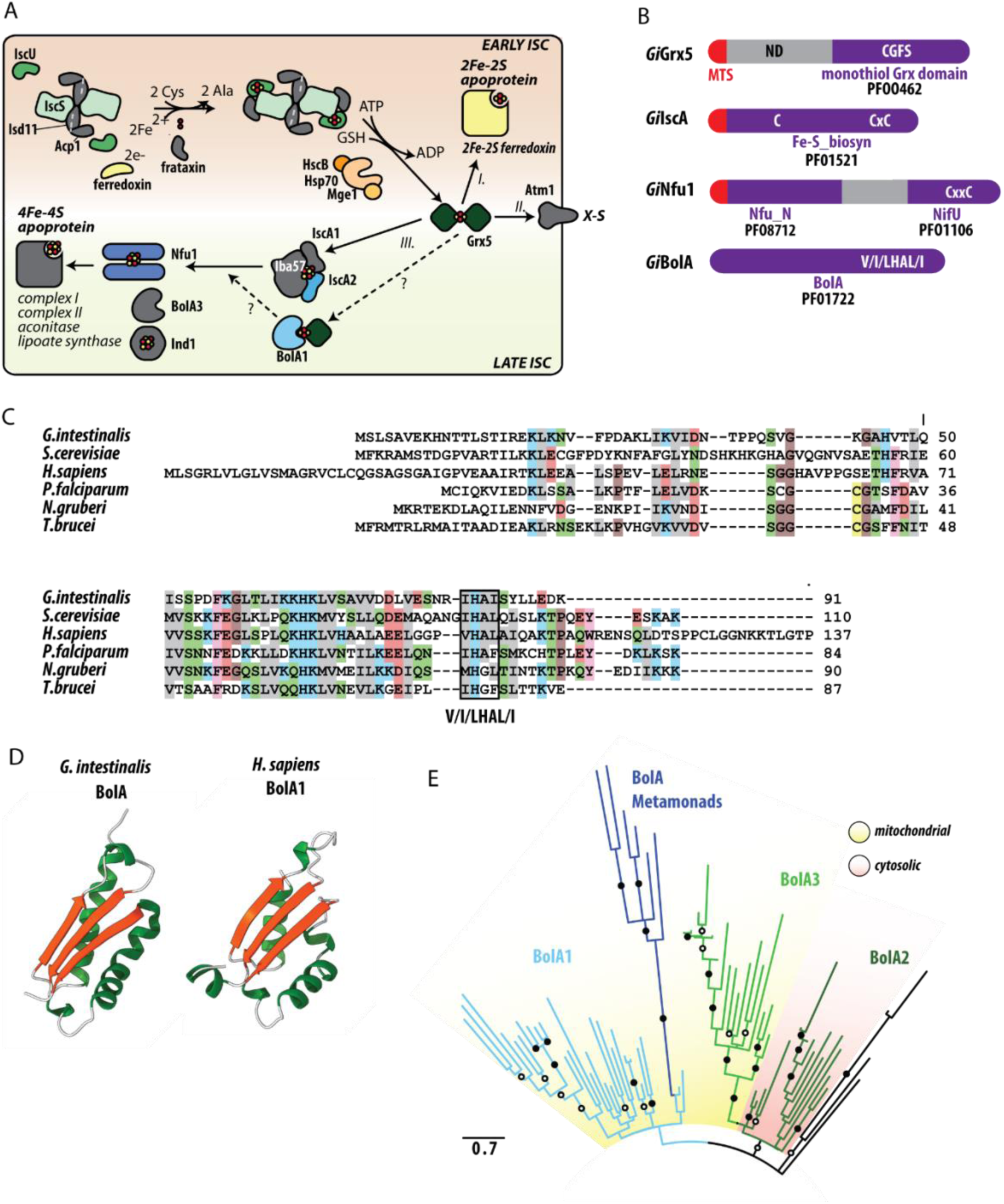
Components of late ISC pathway in *G. intestinalis*. (A) Schematic representation of mitosomal ISC pathway. The mitochondrial components that are missing in *Giardia* mitosomes are shown in grey. Early and late ISC pathway is distinguished by the background colour, [2Fe-2S] cluster on Grx5 dimer can be (*I.*) transferred to the target mitochondrial apoproteins (*II.*) exported to the cytosol or (*III.*) enters the late ISC machinery. (B) Domain structure of *Gi*Grx5, *Gi*IscA, *Gi*Nfu1, and *Gi*BolA. The respective sequence motifs and Pfam accession numbers are shown. (C) Protein sequence alignment of the identified *Gi*BolA with the homologues from, *Saccharomyces cerevisiae* (Q3E793), *Homo sapiens* (Q9Y3E2), *Plasmodium falciparum* (Q8I3V0), *Naegleria gruberi* (D2V472) and *Trypanosoma brucei* (Q57YM0). BolA signature V/I/LHAL/I motif is highlighted. (D) Structure of *Gi*BolA as predicted by AlphaFold2 (73), predicted structure of human BolA1 (*Hs*BolA1) (27) is shown for comparison. (E) Maximum likelihood phylogenetic tree of 70 eukaryotic BolA1 paralogues shows that *Gi*BolA and metamonad BolA homologues emerge from within a clade of mitochondrial BolA1 proteins. Summary of bipartition support values (1000 ultrafast bootstraps) greater than 80 or 95 are shown in open and closed circles, respectively.

Grx5 acts as the central dividing point between the early and late ISC pathway at which the assembled [2Fe-2S] cluster is either (i) transferred to the target mitochondrial [2Fe- 2S] apoproteins, (ii) exported to the cytosol as an enigmatic X-S compound or (iii) enters the late ISC machinery (9, 21). The late ISC machinery starts with the transfer of two [2Fe-2S] clusters from Grx5 to a complex of IscA1, IscA2 and Iba57 (22) where the [4Fe-4S] cluster is formed (23). The newly created [4Fe-4S] clusters are delivered to apoproteins with the help of Nfu1 (24, 25) and Ind1, the latter being specifically involved in [4Fe-4S] cluster-binding for the complex I assembly (26). Recently, two conserved factors BolA1 and BolA3 have been shown to participate in the transfer of [4Fe-4S] clusters to apoproteins in mitochondria (27). BolA1 and BolA3 have overlapping functions, but preferentially act on Grx5 and Nfu1, respectively (25). Importantly, BolA function has previously been associated with aerobic metabolism, which was supported by its absence in anaerobic eukaryotes (28).

It is now generally accepted that the early ISC pathway is a converging evolutionary point of the MROs, *i.e*., no matter how much the mitochondrion has been modified during evolution, most MROs have retained early ISC components like IscU and IscS (29). Moreover, some of the MROs like mitosomes of *G. intestinalis* also contain components of the late ISC pathway.

Therefore, here, we sought to experimentally examine the nature of the late ISC pathway in *G. intestinalis.* Using enzymatic tagging and series of affinity pulldowns, we have generated a robust interactome of the mitosomal late ISC pathway revealing that Grx5, Nfu1 and herein discovered BolA orthologue are at the core of the pathway. On the other hand, mitosomal IscA appears to function in downstream steps of the pathway. The specific interaction between BolA and Grx5 could be confirmed by yeast two hybrid assays as well as by their strict co-occurrence in other MRO-carrying species. However, no endogenous mitosomal substrate for the late ISC pathway could be identified in the mitosomal proteome or in the bioinformatic search of the *G. intestinalis* genome. Hence, a complete *bolA* knockout strain was generated by CRISPR/Cas9 which showed a significantly decreased activity of cytosolic [4Fe-4S] pyruvate:ferredoxin oxidoreductase. These results indicate that mitosomal BolA, and thus the late ISC pathway, is required for the formation of cytosolic [4Fe-4S] clusters. Such functional connection is unknown for mitochondria and may represent unique of adaptation of MROs.

## RESULTS

### The late ISC pathway and the identification of BolA in *G. intestinalis*

Previous genomic and proteomic analyses of *G. intestinalis* revealed the presence of three late ISC pathway components; Nfu1, IscA and Grx5, hereafter referred to as *Gi*Nfu1, *Gi*IscA, and *Gi*Grx5, respectively (Fig. 1A). All three proteins possess highly conserved cysteine residues that are necessary for the coordination of the Fe-S cluster. *Gi*Grx5 contains the CGFS motif of monothiol glutaredoxins (Fig. 1B, Supplementary Fig. 1A), the C-terminal domain of *Gi*Nfu1 carries a CxxC motif (Fig. 1B and Supplementary Fig. 1B) and *Gi*IscA carries a CxnCxC signature motif (Fig. 1B, Supplementary Fig. 1C). Both *Gi*Nfu1 and *Gi*IscA carry a short N-terminal pre-sequence that likely serves as the mitosomal targeting signal. *Gi*Grx5 was previously shown to carry a long non-homologous N-terminal sequence, which is required for targeting but may possibly play an additional role in protein function (30). Of the two types of IscA proteins known for eukaryotes, only IscA2 was identified in *G. intestinalis* (4).

The presence of these three late ISC components in *G. intestinalis* prompted us to search for other factors that were identified within the late pathway. Specifically, the orthologues of BolA, Iba57 and Ind1 proteins were searched using hidden Markov model (HMM) profiles against the *G. intestinalis* genome. Interestingly, while the last two searches did not result in the identification of positive hits, a single BolA orthologue was identified in *G. intestinalis* (*Gi*BolA) (Fig. 1B, 1C). The protein could be readily identified in the conceptual proteomes of all genotypes (assemblages) including new genome assembly of WBc6 (31) but was missing from the original reference genome, probably due to its small size (32). The amino acid sequence of *Gi*BolA contains signature V/I/LHAL/I motif towards the C-terminus (33) but no putative N-terminal targeting sequence, as is common to most other BolA proteins (e.g., Fig. 1C). Structural prediction of *Gi*BolA using AlphaFold 2 revealed an αβαβ topology that matches experimentally solved or predicted structures of BolA homologs from both eukaryotes and prokaryotes (Fig. 1D) (34, 35). The only structural difference is a short C- terminal α-helix missing in *Gi*BolA (Fig. 1D). Given the occurrence of three BolA proteins in eukaryotes, phylogenetic analysis was performed to determine which of three eukaryotic BolA paralogues, functioning in the cytosol (BolA2) (36) or mitochondria (BolA1 and BolA3) (28, 37) is present in *G. intestinalis.* The analysis showed that *Gi*BolA and other BolA proteins that could be identified in the Metamonada supergroup emerge from within a clade of BolA1 proteins (Fig. 1E) suggesting that *G. intestinalis* contains an orthologue of mitochondrial BolA1, which would hence be expected to be localized in mitosomes.

### *Gi*BolA is part of mitosomal late ISC pathway

To test whether *Gi*BolA is indeed a mitosomal component, the protein was expressed with the C-terminal biotin acceptor peptide tag (BAP) tag. Immunodetection of the tag by fluorescence microscopy showed clear colocalization of *Gi*BolA with the mitosomal marker GL50803_9296 (Fig. 2A). Western blot analysis of the cellular fractions revealed the specific presence of the protein in the high-speed pellet (HSP) fraction that is enriched for mitosomes (Fig. 2B). Except for *Gi*Grx5 (30), the mitosomal localization of other late ISC components had not been previously experimentally confirmed. Therefore, analogously, all three proteins were expressed with the C-terminal BAP tag and their cellular localization was detected in the fixed cells (Fig.2A) and in the cell fractions (Fig. 2B). All proteins specifically localize in the mitosomes. Furthermore, we tested whether BAP-tagged proteins are localized within the mitosomes or are accumulated on the surface of the organelle as a possible result of protein overexpression. To this end, a protease protection assay was performed on *G. intestinalis* expressing BAP-tagged proteins whereby HSPs were incubated with trypsin in presence or absence of a membrane-solubilizing detergent. Proteins encased by one or more membranes will be inaccessible to trypsin and will therefore be detected by standard immunoblotting in the absence but not presence of the detergent (Fig. 2C). Unlike the outer membrane marker *Gi*Tom40, all late ISC components were resistant to protease treatment as the mitosomal matrix marker IscU. As a control, mitosomal membrane solubilization resulted in overall protein degradation. In summary, all four proteins were found specifically located within mitosomes, suggesting that the minimalist late ISC pathway occurs within the organelles.

**Figure 2.**
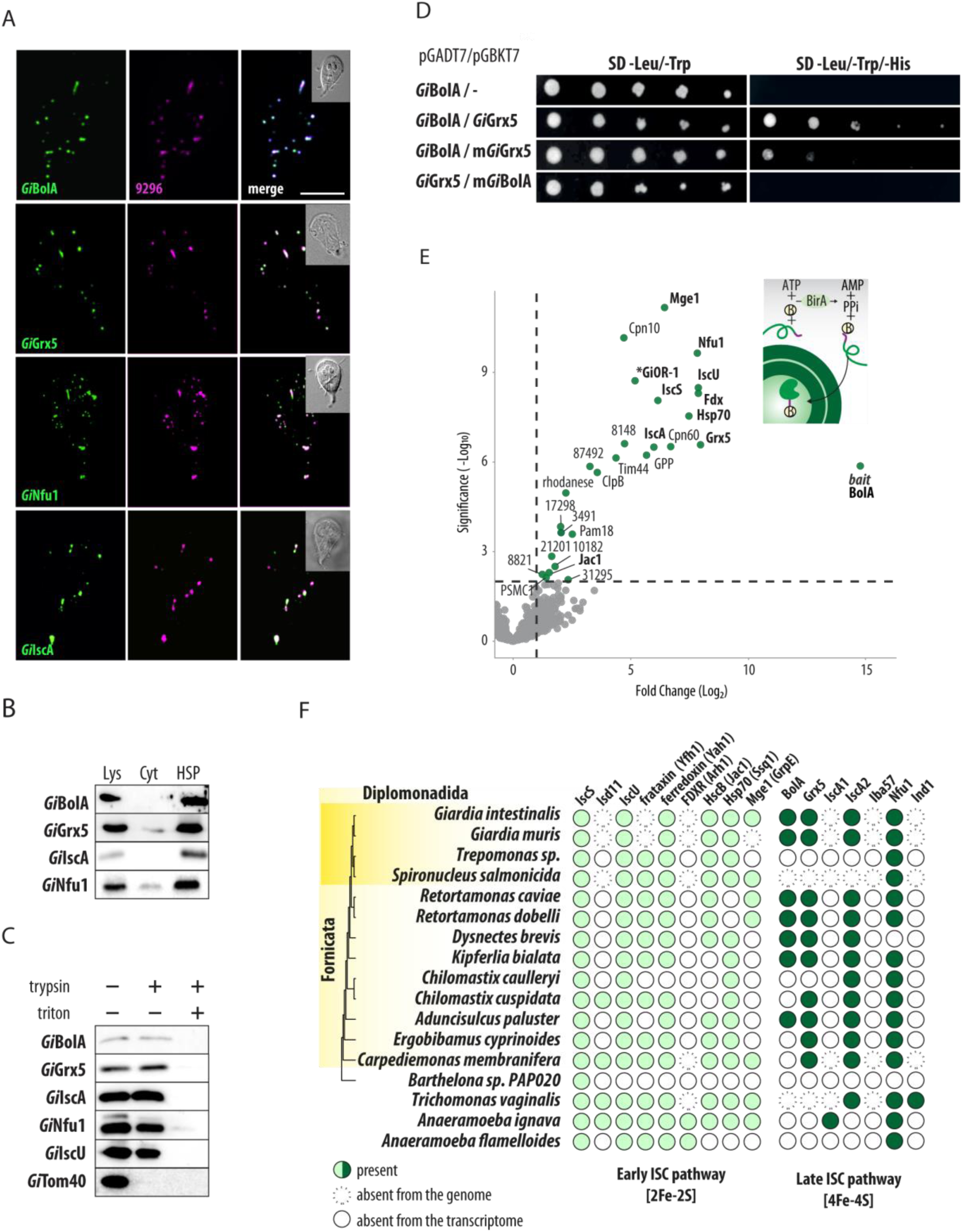
*Gi*BolA is a mitosomal protein that specifically interacts with *Gi*Grx5 and other ISC components. (A) BAP-tagged *Gi*BolA, *Gi*Grx5, *Gi*Nfu1 and *Gi*IscA were expressed in *G. intestinalis* and the proteins were detected by anti-BAP antibody (green). The co-colocalization with mitosomal marker GL50803_9296 (magenta) is shown. The DIC image of the cell is shown in the inlet, the scale bar represents 5 μm. (B) Detection of BAP- tagged *Gi*BolA, *Gi*Grx5, *Gi*Nfu1 and *Gi*IscA in cellular fractions, lys – cell lysate, cyt - cytosol, HSP – high speed pellet fraction. (C) Protease protection assay of late ISC components and the markers of the outer mitosomal membrane (*Gi*Tom40) and the mitosomal matrix (*Gi*IscU). High-speed pellets isolated from *G. intestinalis* expressing BAP-tagged *Gi*BolA, *Gi*Grx5 *Gi*IscA and *Gi*Nfu1 were incubated with 20 μg/ml trypsin and 0.1% Triton X-100. The samples were immunolabeled with antibodies against the BAP tag, *Gi*Tom40 and *Gi*IscU. (D) Serial dilutions of Y2H assay testing the protein interactions between *Gi*BolA and *Gi*Grx5. The introduction of specific mutations of conserved residues (H90A *Gi*BolA and C128A *Gi*Grx5) abolished the interaction, double and triple dropout medium was used to test the presence of the plasmids and the interaction of the encoded proteins, respectively. (E) Affinity purification of the *in vivo* biotinylated *Gi*BolA with the DSP-crosslinked interacting partners. (top right) Scheme of the *in vivo* biotinylation of the C-terminal BAP-tag of *Gi*BolA by cytosolic BirA. (left) Volcano plot of the statistically significant hits obtained from the protein purification on streptavidin coupled Dynabeads. Components involved in ISC pathway are shown in bold letters. (F) The presence/absence of the ISC components in Metamonada supergroup.

### Mitosomal BolA specifically interacts with Grx5 and other mitosomal ISC components

Recent studies on human BolA proteins showed a specific interaction of mitochondrial BolA1 with Grx5 during the stabilization of [2Fe-2S] cluster on Grx5 (27). Using a yeast two hybrid (Y2H) assay, we tested whether mitosomal BolA also interacts with Grx5. Indeed, the assay was able to show the interaction between *Gi*BolA and *Gi*Grx5 (Fig. 2D). Previous studies in yeast identified the specific residues of BolA and Grx5 critical for interaction (27). Therefore, we tested whether the same molecular interaction can also be demonstrated for the *Giardia* proteins. Specifically, the cysteine residue (position 128) within the CGFS motif of *Gi*Grx5 and a highly conserved histidine residue (position 82) of *Gi*BolA, that were both shown to coordinate Fe-S cluster (38). In both cases, the introduced mutations abolished the positive interaction in Y2H (Fig. 2D). These results strongly suggest that the mechanism of interaction is conserved for the late ISC components in the *G. intestinalis* mitosomes. However, the analogous assay did not show any interaction between *Gi*BolA and *Gi*Nfu1 (data not shown), that would be expected if *Gi*BolA represented a BolA3 homologue (25)

To reveal the complex *in vivo* interactions of *Gi*BolA, we used a previously established method of enzymatic tagging in *G. intestinalis* that is based on co-expression of the biotin ligase (BirA) and protein of interest tagged by BAP (3). In the presence of ATP, BirA specifically biotinylates the lysine residue within the BAP tag. Therefore, a BAP-tagged *Gi*BolA was introduced into *G. intestinalis* expressing cytosolic BirA. The mitosomes- enriched HSP was incubated with the chemical crosslinker DSP and *Gi*BolA-BAP was purified on streptavidin-coupled magnetic beads (see Materials and Methods for more details). The purified crosslinked complexes were subjected to proteomic analysis and the resulting peptide mass spectra were searched against the predicted proteome of *G. intestinalis* (39). Data obtained from the biological and technical triplicates (Supplementary Table 1) were displayed in a volcano plot showing the fold change of protein abundance compared to the negative control (Fig. 2E). In total, 26 significantly-enriched proteins were identified. *Gi*Grx5 represented the most enriched interactor but other ISC components (NifU, IscA, Fdx, IscU, IscS, Hsp70, Jac1) also appeared among the most significant enriched proteins (Fig. 2E). The remaining proteins represented mitosomal proteins involved in protein import and folding, and mitosomal proteins of unknown function. At least one probable non-mitosomal protein (PSMC1, Proteasome 26S Subunit, ATPase 1 homologue) was identified among the significantly enriched proteins (Fig. 2E, Supplementary Table 1) suggesting minimal contamination from non-mitosomal proteins in this proximity tagging method. The dominant presence of mitosomal matrix proteins in the presented interactome strongly suggests that *Gi*BolA is localized in the mitosomal matrix. This represents the first report of a BolA protein and putative late ISC pathway in an anaerobic mitochondrial organelle.

### Co-occurrence of mitochondrial BolA and Grx5 and a uniform pattern of late ISC components in metamonads

All known eukaryotic organisms belonging to the Metamonada supergroup of eukaryotes carry MROs adapted to life without oxygen. According to genomic and transcriptomic analyses, the degree of metabolic reduction of these MROs varies across the Metamonada (40, 41). Some MROs participate in ATP generation and some, such as *G. intestinalis* mitosomes, are involved only in the synthesis of Fe-S clusters. The identification of *Gi*BolA prompted us to search the available data for the homologues of BolA and other ISC components in Metamonada.

A BolA homologue was detected in genomes of the parasitic *Giardia muris* and two *Retortamonas* species, and in free-living *Dysnectes brevis*, *Kipferlia bialata* and *Aduncisulcus paluaster* (Fig. 2F, Supplementary Table 2). Similarly to *G. intestinalis*, the vast majority of Metamonada have been found to lack Iba57 and IscA1. The absence of the former correlates with the absence of complex I in these eukaryotes, but both Iba57 and IscA1 are supposed to constitute a complex together with IscA2, on which the [4Fe-4S] cluster is formed (42) This raises the general question whether IscA2, unlike the whole IscA1-IscA2-Iba57 complex, has an indispensable role for anaerobic eukaryotes. Analogously, we could not detect the early ISC components Isd11 and ferredoxin reductase (Arh1) in preaxostylids and fornicates (Fig. 2F). These components were only detected in the less reduced MROs of parabasalids (e.g., *Trichomonas vaginalis*) and in anaeramoebids. Of course, additional components can be identified in the species with incomplete genomic data, yet these results likely demonstrate the ancestral adaptation of the late ISC pathway in Metamonada that involved the loss of Iba57 and IscA proteins.

### Interactome of late ISC components reveals a downstream role of IscA

Characterization of late ISC pathway in mitochondria has relied largely on genetic and biochemical approaches *e.g.*, (25–27,43–45). Here, we chose to continue with the affinity-purification proteomics, which to our knowledge has not yet been used in this context, to characterize the pathway in *G. intestinalis* mitosomes. The combination of protein specific interactomes as the one obtained above for *Gi*BolA can yield a spatial reconstruction of the pathway (46). In addition, it can also identify putative mitosomal client apoprotein(s) that receive the synthesized [4Fe-4S] clusters as it was done for its mitochondrial counterparts (25, 47). To this aim, proteins co-purified in complexes chemically crosslinked to *Gi*Grx5, *Gi*Nfu1, and *Gi*IscA were identified by mass spectrometry. The returned datasets contained 47, 30, and 22 statistically significant proteins of three independent sets of experiments, respectively (Fig. 3A-C, Supplementary Table 1).

**Figure 3.**
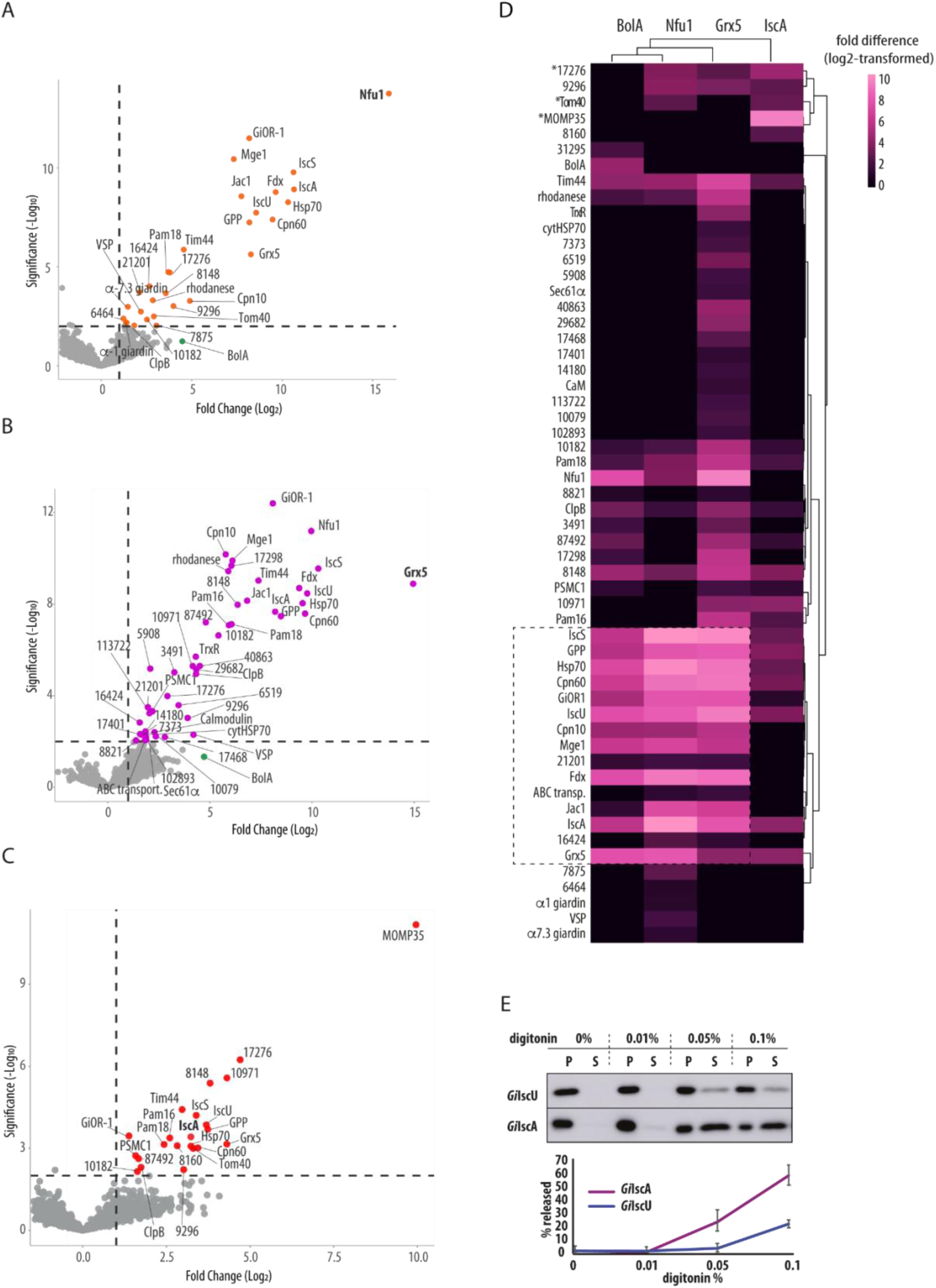
Proteomic analysis of late ISC pathway. BAP-tagged *Gi*Grx5, *Gi*Nfu1 and *Gi*IscA were *in vivo* biotinylated by cytosolic BirA and purified on streptavidin-coupled Dynabeads upon crosslinking by DSP. (A-C) Volcano plots depict the significantly enriched proteins that co-purified with (A) *Gi*Nfu1, (B) *Gi*Grx5 and (C) *Gi*IscA. (D) Heatmap of combined significantly enriched proteins for all four late ISC components, (E) Digitonin solubilization of the mitosomes shows differential release of IscA over IscU, P -pellet fraction (retained protein), S – supernatant (released protein). Exemplary western blot of four independent experiments is shown, the error bars show standard deviation.

The final combined dataset which also included the *Gi*BolA purification data was plotted in a heat map using log2 transformed fold difference values (Fig. 3D). Hierarchical clustering showed a close relationship between the *Gi*BolA-, *Gi*Nfu1- and *Gi*Grx5-specific protein profiles, while the *Gi*IscA-specific dataset remained the most distinct. The interactomes of the first three proteins converged over the ISC components, chaperones and the mitosomal processing peptidase (GPP) that corresponds to the ‘core’ of the mitosomal metabolism (dashed line in Fig 3D). Several low abundance proteins of unknown function (GL50803_21201, GL50803_16424 and ABC transporter GL50803_87446) were also found in the cluster. Interestingly, a thioredoxin reductase (TrxR) homolog (GL50803_9287) was found among several proteins unique to the *Gi*Grx5 dataset (Fig. 3B). The protein was previously characterized in *G. intestinalis* as cytosolic protein, yet without any interacting thioredoxin (48). Our data suggested that TrxR thus could also act in the mitosomes and reduce *Gi*Grx5 to act as a missing reductase system. *Gi*BolA was found among enriched proteins in *Gi*Grx5 and *Gi*Nfu1 datasets (Fig. 3A, 3B) yet it was not a significant hit due to the incomplete coverage in some of the technical triplicates within biological triplicates. This indicates lower expression levels of *Gi*BolA when compared to other late ISC components.

In contrast, the *Gi*IscA dataset showed enrichment of the outer mitosomal membrane proteins MOMP35 and GL50803_17276 (3, 49). Additionally, Tom40, a central component of the outer membrane translocase, was identified among the significantly enriched proteins (Supplementary table 1). Unlike the interactomes of the other ISC components, many of the ’core’ mitosomal matrix proteins were not significantly enriched in the GiIscA interactome. The affinity of *Gi*IscA to the outer membrane proteins suggested the possibility that the protein is not localized, at least not completely, in the mitosomal matrix but in the intermembrane space (IMS) or it is associated with the outer mitosomal membrane. The latter could be rejected due to the lack of any transmembrane domains and due to the full protection of *Gi*IscA against the externally added protease (Fig. 2C). Therefore, the presence of the protein in the IMS was tested. We took advantage of differential sensitivity of the outer and inner mitosomal membranes to digitonin lysis (3, 50).

The mitosome-enriched fraction was isolated from cells co-expressing *Gi*IscA and the matrix marker *Gi*IscU and incubated with the increasing concentration of digitonin. The release of the proteins from the organelles was monitored via Western blot (Fig. 3E). Interestingly, *Gi*IscA showed a greater proportion of protein released into the supernatant fraction than *Gi*IscU, supporting the hypothesis that *Gi*IscA and *Gi*IscU are not in the same mitosomal subcompartment.

### Mitosomes likely lack the [4Fe-4S] client for the late ISC pathway

The late-acting ISC machinery is responsible for the formation of [4Fe-4S] cluster and its delivery to the client apoproteins within the mitochondrion of model eukaryotes. These include many mitochondrial proteins functioning in the electron transport chain, the TCA cycle, and cofactor biosynthesis (51–54). However, all these proteins are absent in the highly reduced *G. intestinalis* mitosomes.

To identify possible mitosomal clients of the late ISC pathway, we first investigated the interactome of *Gi*Nfu1 with the premise that the apoproteins that receive their [4Fe-4S] cluster from the late ISC pathway can be co-purified with Nfu1 (25,47,55). The search in the dataset for [4Fe-4S] cluster motifs (56) did not return any positive hits, therefore an unbiased search for Fe-S proteins within the entire conceptual *G. intestinalis* cellular proteome was performed by MetalPredator (57). Upon manual checking with available literature and structural information, 40 proteins were identified that bind [4Fe-4S] clusters (Fig. 4A, Supplementary Table 3). Of these, 19 were predicted to function in the cytosol in energy, redox, amino acid, and nitrogen metabolism, as well as cofactor biosynthesis and protein translation. There were 11 nuclear proteins identified, participating either in DNA or RNA metabolism. The remaining components corresponded to the transient cluster carriers of the mitosomal ISC machinery and cytosolic iron–sulfur assembly (CIA) pathway (58). The only mitosomal protein with stably associated Fe-S cluster is [2Fe-2S] ferredoxin, which is itself directly involved in the ISC pathway as an electron carrier. Of course, we cannot rule out the presence of a previously unknown protein with a unique cluster binding domain/motif in mitosomes, but the present data suggest that mitosomes lack any client [4Fe-4S] protein for their late ISC pathway.

**Figure 4.**
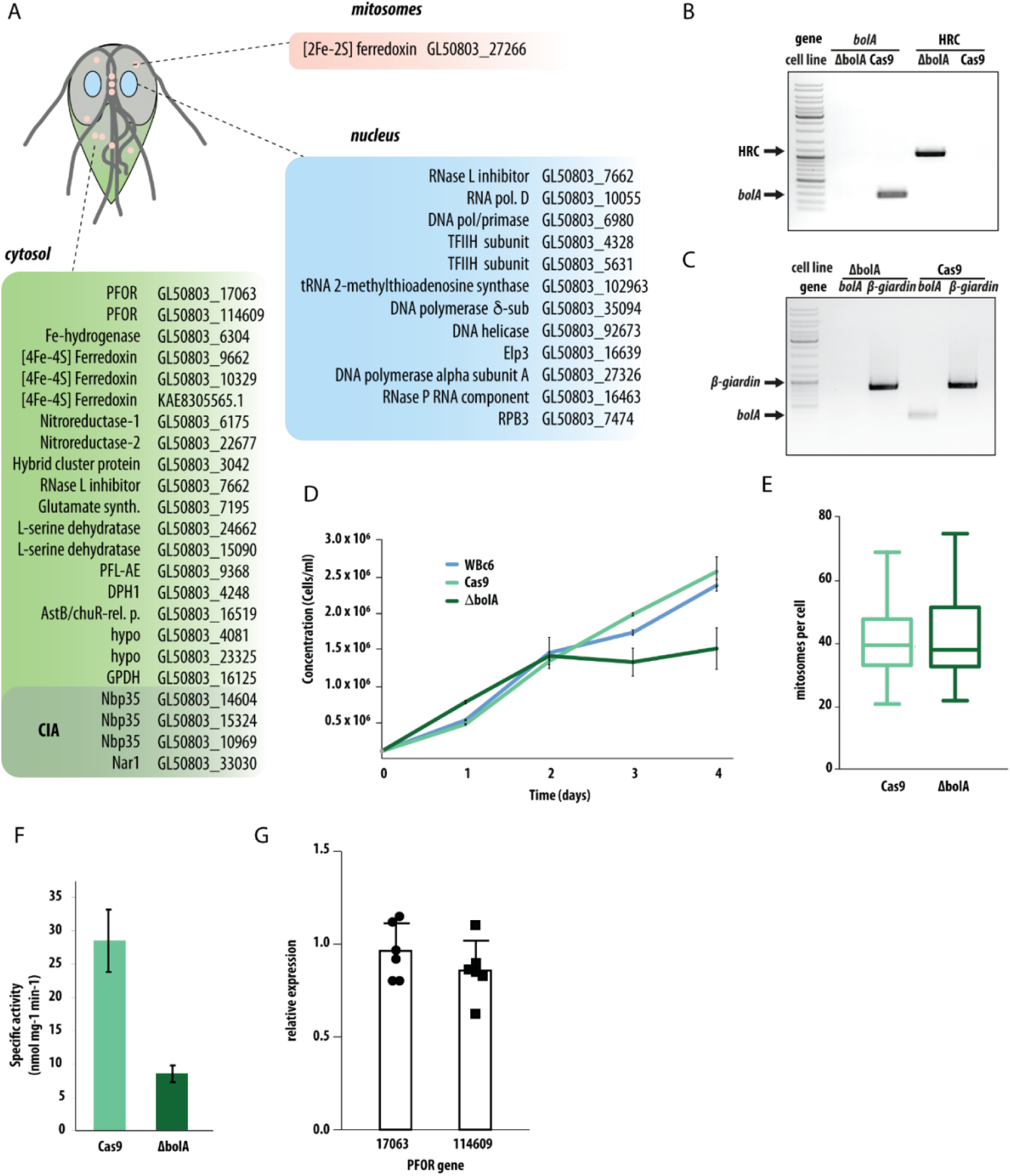
Mitosomal *Gi*BolA is involved in the formation of cytosolic Fe-S proteins. (A) The list of predicted 40 Fe-S proteins in *G. intestinalis* includes only one mitosomal protein, [2Fe-2S] ferredoxin, that itself participates in the ISC pathway. All putative clients that require [4Fe-4S] clusters are localized in the cytosol or in the nucleus (Supplementary Table 3). (B) The ΔbolA cell line was tested for the presence of *bolA* gene and the integration of homologous recombination cassette (HRC) by PCR on gDNA, (C) the expression of *bola* gene in ΔbolA cell line was tested by PCR on the cDNA, *β-giardin* was used as a control gene. (D) The slowed growth phenotype of ΔbolA cell line in comparison to parental Cas9- expressing cell line and wildtype WBc6 strain, error bars represent standard deviation. (E) The number of mitosomes per cells in Cas9-expressing (n=64) and ΔbolA cells (n=107), the error bars of the box plot depict min to max values. (F) Decreased activity of cytosolic PFOR in ΔbolA cell line when compared to the parental Cas9-expressing cell line, the error bars depict standard deviation. (F) Real-time PCR results show relative expression of two PFOR-encoding genes, GL50803_17063 and GL50803_114609, in ΔbolA cell line. Expression levels are depicted relative to the control cell line. Calculated results from six independent RNA isolations are shown for each gene. The expression of both genes was normalized to NADH oxidase-encoding gene, GL50803_33796. Cas9-expressing cell line was used as a control, the error bars depict standard deviation.

### Knockout of *bolA* gene is manifested by a decrease in the activity of the cytosolic [4Fe- 4S] containing PFOR

BolA was previously thought to be restricted to aerobic eukaryotes (28), thereby all functional analyses have been performed on aerobic model organisms (59). Having established the integration of *Gi*BolA within the mitosomal late ISC pathway, we next examined the role of BolA in the formation of Fe-S clusters. To this aim, using the recently established CRISPR/Cas9-mediated gene knockout approach (60) and a *G. intestinalis* cell line lacking *bolA* gene (ΔbolA) was generated (Fig. 4B, 4C). The gene knockout was verified by PCR on the gDNA for the absence of *bolA* gene and the presence of homologous recombination cassette (HRC) (Fig. 4B). Furthermore, no *bolA* mRNA was detected in cDNA prepared from the cells (Fig. 4C). Finally, the proteomic analysis of mitosomes-enriched HSP fraction showed the absence of BolA when compared to the control (Supplementary Table 4).

The ΔbolA cell line exhibited slowed growth when compared to the parental cells (Cas9) and the wild-type control (WBc6) (Fig. 4D) but the overall number and morphology of the mitosomes remained unchanged (Fig. 4E, Supplementary Fig. 2). This indicated that disruption of the function of late ISC pathway can perturb growth rate but not mitosomal morphology or number in *G. intestinalis*. Given the apparent absence of the client proteins in the mitosomes, the formation of [4Fe-4S] clusters was monitored indirectly via the activity of cytosolic enzyme pyruvate-ferredoxin oxidoreductase (PFOR). PFOR catalyses oxidative decarboxylation of pyruvate and produces acetyl-CoA and CO2 with concomitant reduction of cytosolic ferredoxin (another [4Fe-4S]-containing protein), hence acting as cytosolic alternative of pyruvate dehydrogenase complex in mitochondria of aerobes (61). Indeed, the specific activity of PFOR was more than three times lower in ΔbolA cells when compared to the control (Fig. 4F). The expression of two *pfor* genes present in *G. intestinalis* genome was measured by qPCR and found almost unchanged in the ΔbolA cells (Fig. 4G). Taken together, these data strongly suggested that the absence of mitosomal BolA impacts the formation of [4Fe-4S] clusters in *G. intestinalis* cytosol.

## DISCUSSION

This study presents the characterization of late ISC pathway in anaerobic protist *G. intestinalis.* While it shows an unexpected presence of BolA in its mitosomes, it also demonstrates the involvement of the mitosomes in the formation of Fe-S clusters for cytosolic proteins. Thus, this is the first study supporting the long-proposed hypothesis of MROs as evolutionarily conserved compartments dedicated to control Fe-S cluster biogenesis. Moreover, it shows that, unlike in mitochondria, the defect in the late ISC pathway of *G. intestinalis* mitosomes affects the activity of Fe-S proteins outside the organelle.

The independent evolution of mitochondria in various anaerobic lineages of eukaryotes resulted into remarkably uniform metabolic adaptations. Comparative studies on mitochondria and various MROs have suggested that the mitochondrial formation of Fe-S clusters was the main selection pressure for retaining the organelles even in the anoxic environments (5,62–66). Mitochondria initiate the biosynthesis of cellular Fe-S clusters via the action of early ISC components that results into the formation of [2Fe-2S] cluster bound by glutaredoxin (Grx5) dimer. From here, the cluster is either distributed to mitochondrial clients, combined via late ISC components to [4Fe-4S] clusters or exported as an unknown sulfur-containing factor to the cytosol (63). Most of mitochondrial Fe-S client proteins contain [4Fe-4S] clusters and thus the late ISC pathway is vital for the function of the respiratory chain, the TCA cycle as well as the synthesis of prosthetic groups such as heme, lipoic acid or molybdenum cofactor (63). Number of late ISC components are dedicated to serve these multiple clients in mitochondria and some of them could be also identified in *G. intestinalis*.

In this study, we show that despite the loss of all mitochondrial pathways that require the presence of [4Fe-4S] clusters, mitosomes of *G. intestinalis* contain four late ISC components; Grx5, IscA, Nfu1 and the newly identified BolA homologue. In classical experimental models of yeast and mammalian mitochondria, defective late ISC pathway is often lethal for the cell or at least lead to severe diseases in humans due to multifactorial deficiencies caused in the mitochondrial metabolism (67, 68). In this context, mitosomes represent a unique biological model to study the non-mitochondrial role of the ISC pathway without the interference with mitochondrial metabolism.

Eukaryotes have three BolA proteins that function together with glutaredoxins in chaperoning the Fe-S cluster both in cytosol and mitochondria (25,27,69). Yet, the previous absence of BolA proteins in the anaerobic eukaryotes that carry MROs suggested that BolA proteins are involved in the aerobic metabolism by controlling thiol redox potential (28). Mitochondrial BolA1 and BolA3 were proposed to function as [4Fe-4S] assembly cluster factors via the interaction with Grx5 and Nfu1, respectively (25,27,70). While the BolA3-Nfu1 interaction is required for the final [4Fe-4S] cluster transfer to the apoprotein (Melber et al. 2016), the exact role of BolA1-Grx5 in the preceding steps remains rather unknown. *Gi*BolA specifically interacts with *Gi*Grx5 as demonstrated by Y2H assay and the pulldown experiment. The interaction of *Gi*BolA with *Gi*Nfu1 was not supported by Y2H assay, yet the *Gi*Nfu1 was among the most enriched proteins co-purified with *Gi*BolA. These data indirectly support the results of the phylogenetic reconstructions assigning *Gi*BolA to BolA1 proteins. Interestingly, the search in other anaerobic organisms with MROs revealed co- occurrence of BolA and Grx5 homologues, supporting their mutual interaction in the pathway. However, for yet unknown reason the pair is expendable in some Metamonada species, some of which carry metabolically versatile ATP-producing MROs, *e.g.,* the parabasalids and anaeramoebids.

Based on this data and the existence of BolA deficient yeast cell lines that exhibited relatively mild phenotype (27), the gene was selected for the targeted removal from *G. intestinalis* genome by CRISPR/Cas9. The assumption was that the gene would not be essential for *G. intestinalis* either. In addition, such a viable mutant could also reveal a general function of mitosomes in Fe-S cluster formation. Indeed, removal of the gene encoding *Gi*BolA by CRISPR/Cas9 showed that this protein is not essential for *G. intestinalis* maintained under laboratory conditions. The ΔbolA cell line showed a growth defect that was not associated with reduced mitosomal biogenesis, consistent with the previous observation that *G. intestinalis* does not respond to metabolic perturbations by altering mitosomal dynamics (71). In yeast and patient-derived cell lines, BolA deficiency is manifested by a decrease in the activity of the [4Fe-4S] cluster containing protein succinate dehydrogenase, but also of pyruvate and 2-ketoglutarate dehydrogenases due to impaired lipoylation by [4Fe- 4S] lipoate synthase (27, 37). Since *Giardia* does not encode any of these proteins or any other obvious mitosomal [4Fe-4S] protein, we explored whether BolA deficiency could be manifested in the activity of cytosolic [4Fe-4S] proteins. We found that the cytosolic [4Fe-4S] PFOR had significantly reduced enzymatic activity in cells lacking BolA compared to the control cell line. Transcription of the two *pfor* genes was nearly identical in the ΔbolA cell line, strongly suggesting that the lack of mature [4Fe-4S] cluster in the protein is responsible for the reduced enzymatic activity. These data demonstrate, for the first time, that mitosomes are needed for cytosolic Fe-S cluster biogenesis in *G. intestinalis.* As *Gi*BolA is the first ISC component removed from *G. intestinalis,* it is difficult to assess whether the reduced PFOR activity is a direct consequence of *Gi*BolA deficiency or a broader downstream outcome of a defect in Fe-S cluster formation.

In model aerobic eukaryotes, there is an additional cytosolic acting BolA protein (BolA2) and glutaredoxin (Grx??? Is this known?) that act as chaperones for cytosolic Fe-S clusters. It is thus possible that the single BolA protein of *G. intestinalis* also effects cytosolic Fe-S clusters. However, further studies are needed to understand the actual connection between the mitosomal late ISC pathway and Fe-S proteins in other cellular compartments. However, it is tempting to speculate that similar connection may exist in the aerobes but has remained unrecognized due the crucial role of the ISC pathway for the mitochondria themselves.

In mitochondria, the Atm1 transporter in the inner membrane was shown to link the early ISC pathway with the cytosolic iron–sulphur assembly (CIA) via the transport of an unknown sulphur-containing molecule (72). Atm1 homologue is missing in *G. intestinalis* and so are any other metabolic transporters or carriers. Thus, surprisingly, *Gi*IscA might have a compensatory role as a candidate for the connection between the cytosolic CIA (58) and mitosomal ISC machinery. The specific interaction of *Gi*IscA with the proteins in the outer mitosomal membrane and the sensitivity to the outer membrane solubilization indicated that it may in fact reside, at least partially, in the IMS of the mitosomes. Although such localization would represent a unique adaptation of *G. intestinalis* mitosomes, it would also correspond to the loss of client proteins in these organelles. Of course, further experiments are needed to describe the place of action of *Gi*IscA but the obvious complication is the size of the mitosomes and the lack of any IMS markers.

To conclude, this work shows how late ISC pathway has undergone specific functional adaptations in a eukaryote inhabiting anoxic environments. It shows for the first time that the formation of Fe-S clusters within these highly reduced mitochondria has remained functionally important for the cytosolic Fe-S proteins as known for the ‘classical’ aerobic mitochondria.

## MATERIALS AND METHODS

### Bioinformatics

The structural models of human and *G. intestinalis* BolA were computed using the Google Colab interface of AlphaFold2 (https://colab.research.google.com/github/sokrypton/ColabFold/blob/main/beta/AlphaFold2_a dvanced.ipynb) (73). The multiple sequence alignment was generated with the jackhmmer option. The best scoring structure according to the plDDT score was subsequently refined with the Amber-Relax option. The [Fe-S] proteins were predicted by Metalpredator (74) using the conceptual proteome of *G. intestinalis* WBc6 strain (giardiadb.org).

### Phylogenetic dataset construction and inferences

Human BolA proteins (NP_001307954.1, NP_001307536.2, NP_997717.2) and Giardia intestinailis BolA-like protein were used as a query against NCBI non-redundant (nr) database to retrieve sequences from select Opishthokonta (Danio renio, Mus musculus, Caenorhabditis elegans Schizosaccharomyces pombe Saccharomyces cerevisiae), select Viridiplantae (Glycine_max Arabidopsis_thaliana Chlamydomonas_reinhardtii Chlorella_variabilis) and non-opisthokonts and non-Viridiplantae (by restricting the database to non-opisthokonts and non-Viridiplantae) with an e-value threshold of 1e^-3^. We also examined the predicted proteomes of metamonads available on EukProt (https://www.biorxiv.org/content/10.1101/2020.06.30.180687v2.abstract) and various sequencing intiatives (40,75,76). The resulting queries were clustered based on sequence identity whereby using cd-hit (77) with a cut-off value of 0.9. Sequences were aligned using mafft (--auto) (78) and ambiguously aligned positions were removed using trimal with ‘-gt 0.5’(79). Phylogenetic inference was performed using IQTREE2 to generate 1000 ultrafast bootstraps (-bb 1000) (80) under the LG+C60+G model of evolution (computed using -mset LG+C20,LG+C10,LG+C60,LG+C30,LG+C40,LG+C50,LG). Trees were visualized using FigTree v1.4 and stylized in Adobe Illustrator. Alignments and tree files are available at figshare (https://figshare.com/s/8fbd1368814dbd11192c reserved DOI:10.6084/m9.figshare.19772155).

### Cloning and protein expression

For the expression of BAP-tagged proteins in *G. intestinalis*, the genes were amplified from genomic DNA and inserted into to pONDRA plasmid encoding the C-terminal BAP tag (81). All the primers and the restriction enzymes used in this study are listed in Supplementary Table 5. Transfection was done as previously described (82) For the *in vivo* biotinylation, the cells expressing BAP-tagged proteins were transfected with a pTG plasmid encoding cytosolic BirA gene from *E. coli* (3). For Y2H assay, genes were amplified from gDNA and subcloned to both pGADT7 and pGBKT7 plasmids. Mutated versions of genes for Y2H assay were commercially synthesized (Genscript).

For CRISPR/Cas9-mediated knockout of *bolA* gene, gRNA sequence ATCAGCTCTCCCGACTTCAA was inserted into gRNA cassette of pTGuide vector using (60) two annealed oligonucleotides (see Supplementary Table 5 for primers and restriction enzymes used). The 999 bp of 5’ and 940 bp 3’ homologous arms surrounding *bolA* gene were inserted into pTGuide vector as the homologous arms for the recombination of the resistance cassette (Supplementary Table 5).

### Real-time PCR

Total RNA from ΔbolA and control cell line was isolated independently six times using NucleoSpinTM RNA isolation kit (Macherey-Nagel) according to manufacturer’s protocol. cDNA prepared from these RNA isolations by KAPA SYBR® FAST One-Step kit (Roche) was analyzed directly by qPCR in Real-time PCR cycler RotorGene 3000 (Qiagen) using Rotor-Gene 6.0 software. qPCRs for each gene were performed in technical triplicates in each RNA isolation for both strains and the mean for each gene from individual RNA isolations was used for further calculations. NADH oxidase-encoding gene, GL50803_33769, was used as a housekeeping gene for normalization.

### Cell culture, fractionation and immunoblot analysis

Trophozoites of *G. intestinalis* strain WB (ATCC 30957) were grown in TYI-S-33 medium (83) supplemented with 10% heat-inactivated bovine serum (PAA laboratories), 0,1% bovine bile and antibiotics. Cells were harvested and fractionated as previously described (3). Cells expressing BAP-tagged *Gi*BolA, *Gi*Grx5, *Gi*Nfu1, and *Gi*IscA were harvested and fractionated as previously described (3) Briefly, the cells were harvested in ice cold phosphate buffered saline (PBS, pH 7.4) by centrifugation at 1,000 × g, 4 °C for 10 min, washed in SM buffer (20 mM MOPS, 250 mM sucrose, pH 7.4), and collected by centrifugation. Cell pellets were resuspended in SM buffer supplemented with protease inhibitors (Roche). Cells were lysed on ice by sonication for 2 min (1 s pulses, 40 % amplitude). The lysate was centrifuged at 2,680 × g, for 20 min at 4 °C to sediment the nuclei, cytoskeleton, and remaining unbroken cells. The supernatant was centrifuged at 180,000 × g, for 30 min at 4 °C. The resulting supernatant corresponded to the cytosolic fraction, and the high-speed pellet (HSP) contained organelles including the mitosomes and the endoplasmic reticulum. The *Gi*Nfu1, *Gi*IscA, *Gi*Grx5 and *Gi*BolA proteins were detected by a rabbit anti-BAP polyclonal antibody (GenScript). Mitosomal *Gi*Tom40 and *Gi*IscU were detected with a specific polyclonal antibody raised in rabbits (84). The primary antibodies were recognized by secondary antibodies conjugated with horseradish peroxidase. The signals were visualized by chemiluminescence using an Amersham Imager 600.

### Immunofluorescence microscopy

*G. intestinalis* trophozoites were fixed and immunolabeled as previously described (71, 85). The C-terminal BAP tag of localized mitosomal proteins was detected by a rabbit anti-BAP polyclonal antibody (GenScript). Mitosomal marker GL50803_9296 was detected by a rabbit anti- GL50803_9296 polyclonal antibody (3). The primary antibodies were detected by secondary antibodies included: Alexa Fluor 594 donkey anti-rabbit IgG (Invitrogen), Alexa Fluor 488 donkey anti-mouse IgG (Invitrogen). Slides were mounted in Vectashield containing DAPI (Vector Laboratories).

Static images were acquired on Leica SP8 FLIM inverted confocal microscope equipped with 405 nm and white light (470-670 nm) lasers and FOV SP8 scanner using HC PL APO CS2 63x/1.4 NA oil-immersion objective. Laser wavelengths and intensities were controlled by a combination of AOTF (Acousto-Optical Tunable Filter) and AOBS (Acousto-Optical Beam Splitter) separately for each channel. Emitting fluorescence was captured by internal spectrally-tunable HyD detectors. Imaging was controlled by the Leica LAS-X software. Images were deconvolved using SVI Huygens software with the CMLE algorithm.

Maximum intensity projections and brightness/contrast corrections were performed in FIJI ImageJ software (86).

### Cross-linking, protein isolation, mass spectrometry (MS)

The HSP (10 mg) isolated from each cell line was collected by centrifugation (30 000 x g, 4°C, 10 min) and resuspend in 1 x PBS supplemented with protease inhibitors (Roche) to protein concentration 1.5 mg/ml. The cross-linker DSP (dithiobis(succinimidyl propionate), ThermoScientific) was added to final 100 µM concentration. The sample was incubated 1 h on ice. Crosslinking was stopped by the addition of 50 mM Tris (pH 8.0) followed by 15 min incubation at RT. The sample was collected by centrifugation (30 000 x g, 10 min, RT) and then resuspended in boiling buffer (50 mM Tris, 1mM EDTA, 1% SDS, pH 7.4) supplemented with protease inhibitors. The sample was then incubated at 80 °C for 10 min, collected by centrifugation and the supernatant was diluted 1/10 in the incubation buffer (50 mM Tris, 150 mM NaCl, 5 mM EDTA, 1% Triton X-100, pH 7.4) supplemented with protease inhibitors. Streptavidin-coupled magnetic beads (50 µL of Dynabeads MyOne Streptavidin C1, Invitrogen) were washed three times in 1 ml of the incubation buffer for 5 min and added to the sample, mixed and incubated for 1 h at room temperature and then incubated overnight with gentle rotation at 4°C. The beads with bound protein were washed three times in the incubation buffer (5 ml) supplemented with 0.1% SDS for 5 min, washed in boiling buffer for 5 min and then washed in the washing buffer (60 mM Tris, 2% SDS, 10% glycerol, 0.1% SDC) for 5 min. Finally, the sample was washed twice in 100 mM TEAB (Triethylammonium bicarbonate, Thermofisher) with 0,1% SDC for 5 min. One tenth of the sample was mixed with SDS-PAGE sample buffer supplemented with 20 mM biotin and incubated in 95°C for 5 min. Experimental controls were tested by immunoblotting and then the sample (dry frozen beads with proteins) was analyzed by mass spectrometry. Control sample was processed in the same way. Each sample was done in triplicate. Beads with bound proteins were submitted to tandem mass spectrometry (MS/MS) analysis as previously described except without the detergent washing steps (82). In brief, captured samples were released from beads by trypsin cleavage. Peptides were separated by reverse phase liquid chromatography and eluted peptides were converted to gas-phase ions by electrospray and analyzed using an Orbitrap (Thermo Scientific, Waltham, MA) followed by Tandem MS to fragment the peptides through a quadropole for final mass detection. Data was analyzed using MaxQuant (version 1.6.3.4) (87) with a false discovery rate (FDR) of 1% for both proteins and peptides and a minimum peptide length of seven amino acids. The Andromeda search engine (88) was used for the MS/MS spectra search against the latest version of the *G. intestinalis* database from EuPathDb (http://eupathdb.org/eupathdb/) and a common contaminant database. Modifications were set as follows: Cystein (unimod nr: 39) as static, and methionoine oxidation (unimod: 1384) and protein N terminus acetylation (unimod: 1) as variable. Data analyses were performed using Perseus 1.6.1.3 (89) and visualized as a volcano plot using the online tool VolcaNoseR (fold change 1,significance threshold 2) (90) and as a heatmap using the online tool ClustVis (91).

### Protease protection and digitonin solubilization assays

For protease protection assay, cells expressing BAP-tagged *Gi*BolA, *Gi*Grx5, *Gi*Nfu1, and *Gi*IscA were harvested and fractionated as described above. The HSP fraction (150 μg) was resuspended in 20 μl of SM buffer and supplemented with protease inhibitors, or 20 μg/ml of trypsin or 20 μg/ml of trypsin and 0.1% Triton X-100. The samples were incubated 30 min at 25 °C and then processed for SDS-PAGE.

For digitonin solubilization assay, 100 µg of HSP fractions isolated from cells co-expressing HA-tagged *Gi*IscU and BAP-tagged *Gi*IscA were incubated for 30 min on ice with 0.01 %, 0.05 %, 0.1 %, digitonin, and without digitonin as a control. The samples were diluted by PBS to 800 µl total volume and collected by centrifugation (30 mins, 180,000 × g, at 4 ° C). The resulting pellets were processed for SDS-PAGE and the supernatants were precipitated by 15 % TCA for 30 min on ice and collected by centrifugation for 30 min at 180,000 × g and 4 ° C, the pellets were washed once with 500 µl of ice-cold acetone, centrifuged as before. The samples were resolved by SDS-PAGE, transferred to nitrocellulose membrane and the protein tags were detected by rabbit anti-BAP antibody (Genscript) and rat anti-HA antibody (Roche). The release to mitosomal proteins was quantified by ImageJ (86).

### Y2H assay

The yeast two-hybrid assay (Y2H) was performed as previously described (92). *S. cerevisiae* cells (strain AH109) were co-transformed with two plasmids (pGADT7, pGBKT7) with the following combinations of genes: *Gi*BolA + *Gi*Grx5, *Gi*BolA + *Gi*mGrx5 (C128A- mutated Grx5), *Gi*Grx5 + *Gi*mBolA (H90A-mutated *Gi*BolA). The empty plasmids were used as negative controls. Co-transformants were selected on double dropout plates SD -Leu/-Trp and triple dropout plates SD -Leu/-Trp/-His. The colonies were grown for four days at 30°C. The positive colonies from triple dropout medium were grown overnight at 30 °C, 200 RPM and then the serial dilution test was performed on double and triple dropout plates.

## ACKNOWLEDGEMENTS

The project was supported by grant from the Czech Science Foundation 20-25417S and project ‘Centre for research of pathogenicity and virulence of parasites’ (No. CZ.02.1.01/0.0/0.0/16_019/0000759) to PD funded by European Regional Development Fund and by a grant from Charles University Grant Agency (project number 1396217) to AM. We acknowledge Imaging Methods Core Facility at BIOCEV, institution supported by the MEYS CR (Large RI Project LM2018129 Czech-BioImaging) and ERDF (project No. CZ.02.1.01/0.0/0.0/18_046/0016045) for their support with obtaining imaging data presented in this paper. CWS was supported in part by European Molecular Biology Organization long-term fellowship (ALTF-997-2015) and the Swedish Research Council (Vetenskapsrådet starting grant 2020-05071).

## SUPPLEMENTARY DATA

**Supplementary Figure 1.**
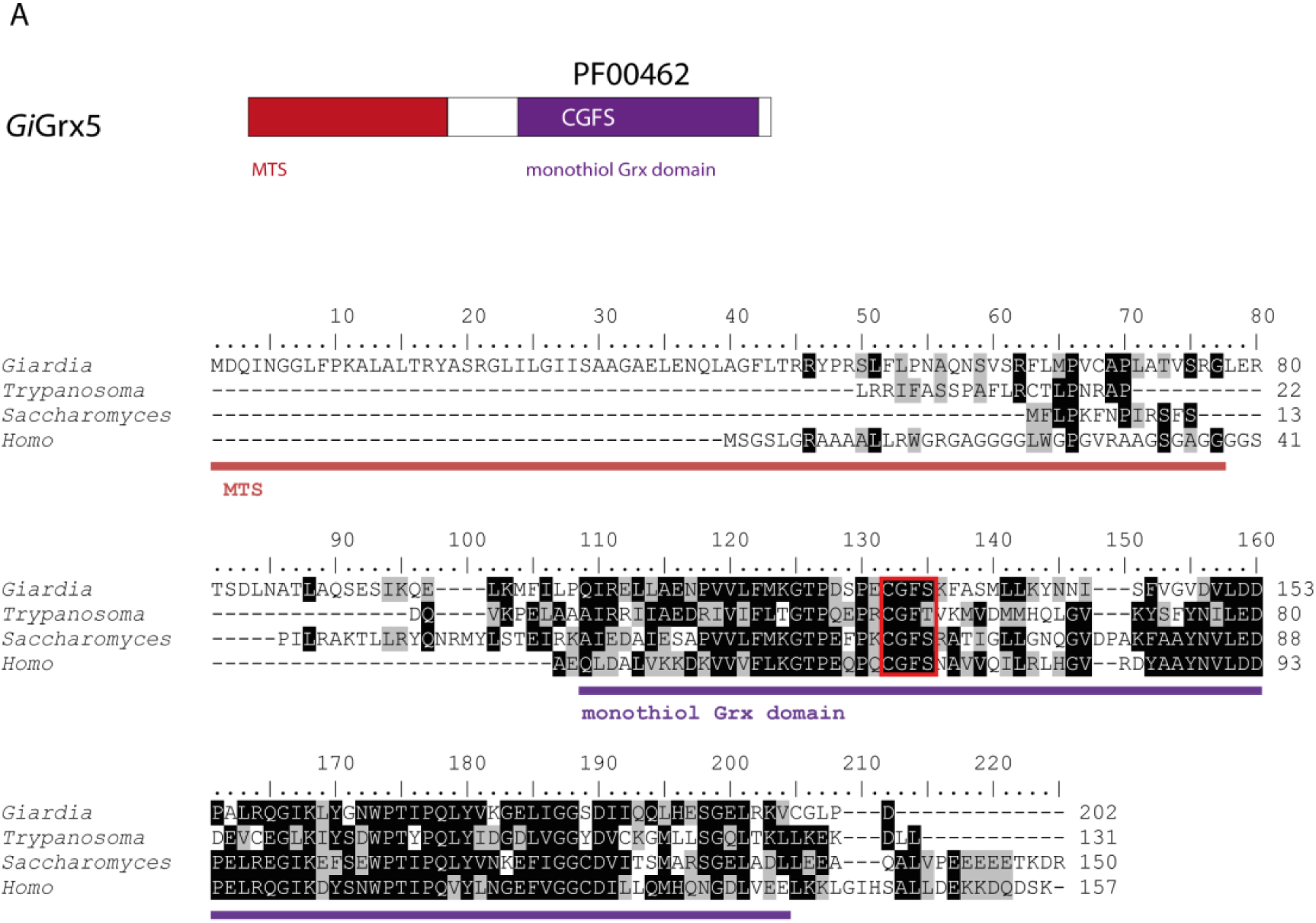
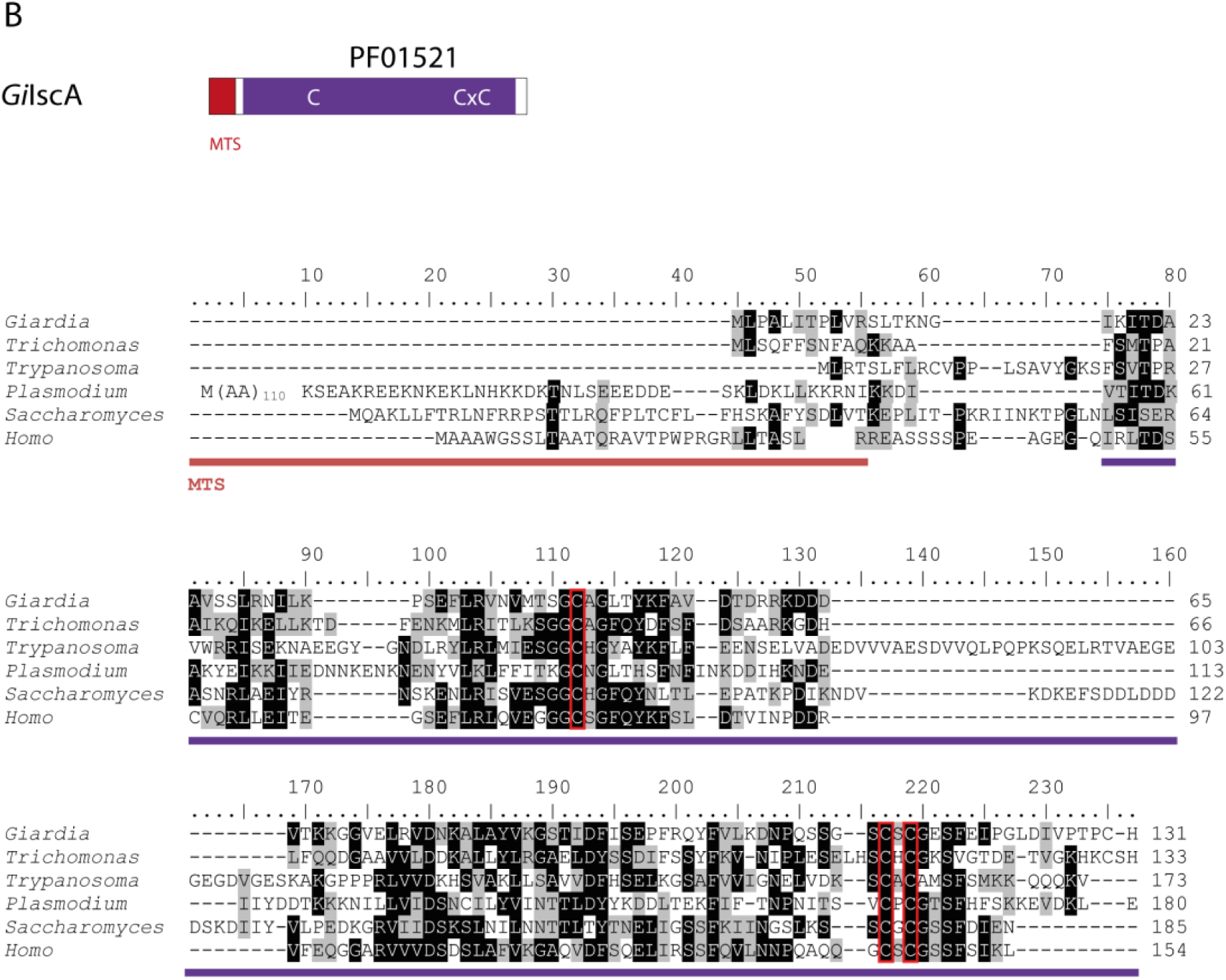
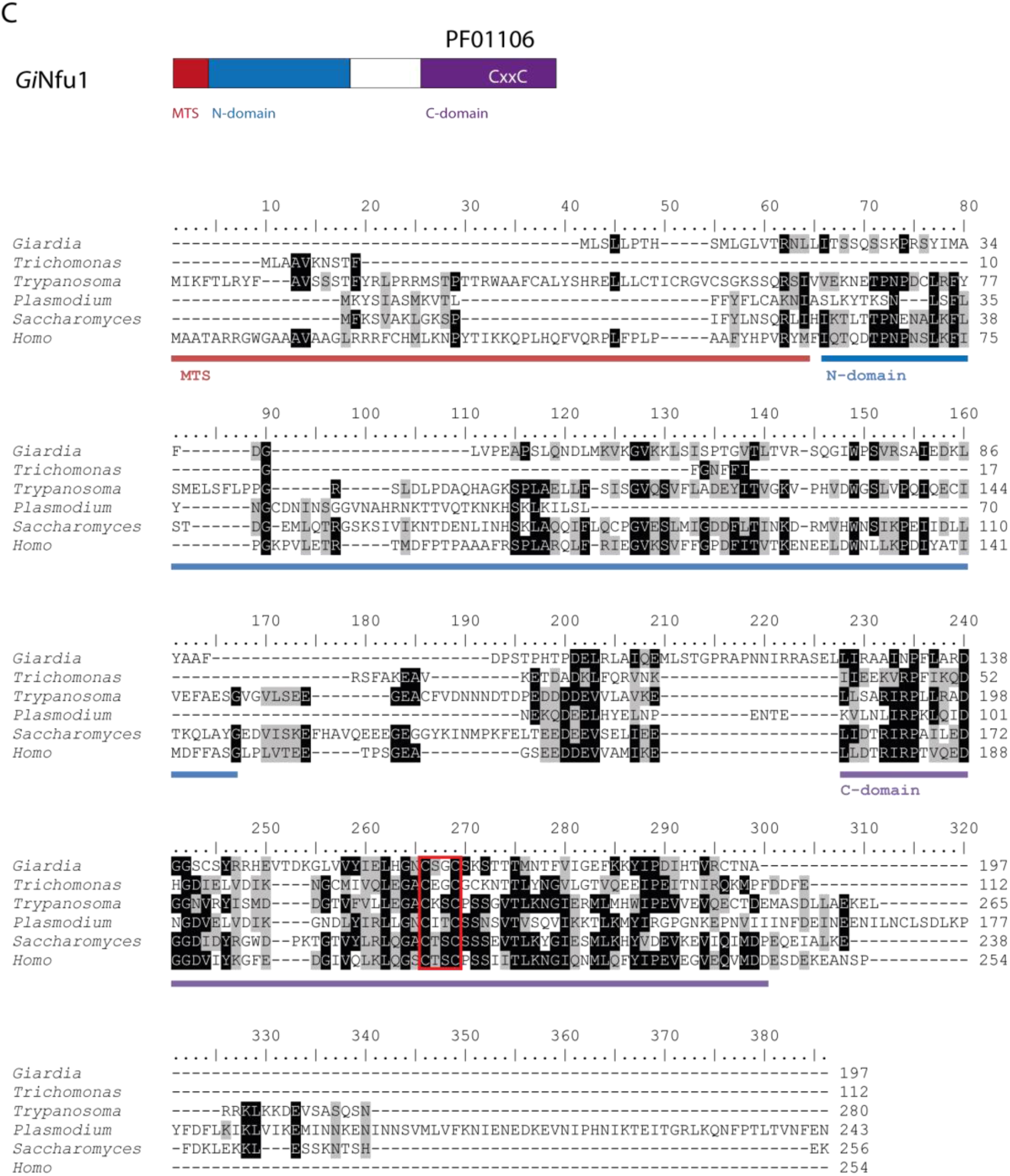
Protein sequence alignments of late ISC components of *Giardia intestinalis.* (A) Grx5, the diagram shows the domain structure of *Gi*Grx5, mitochondrial targeting sequence (MTS) is shown in red, monothiol glutaredoxin domain (PF00462) in purple, the CGFS motif is also highlighted. (B) *Gi*IscA shares the Fe-S_biosyn domain (PF01521) with the conserved cysteine residues involved in cluster binding. (C) *Gi*Nfu1 contains conserved N- and C- domains, the latter of is recognized as NifU domain (PF01106) and carries conserved cysteine motif.

**Supplementary Figure 2.**
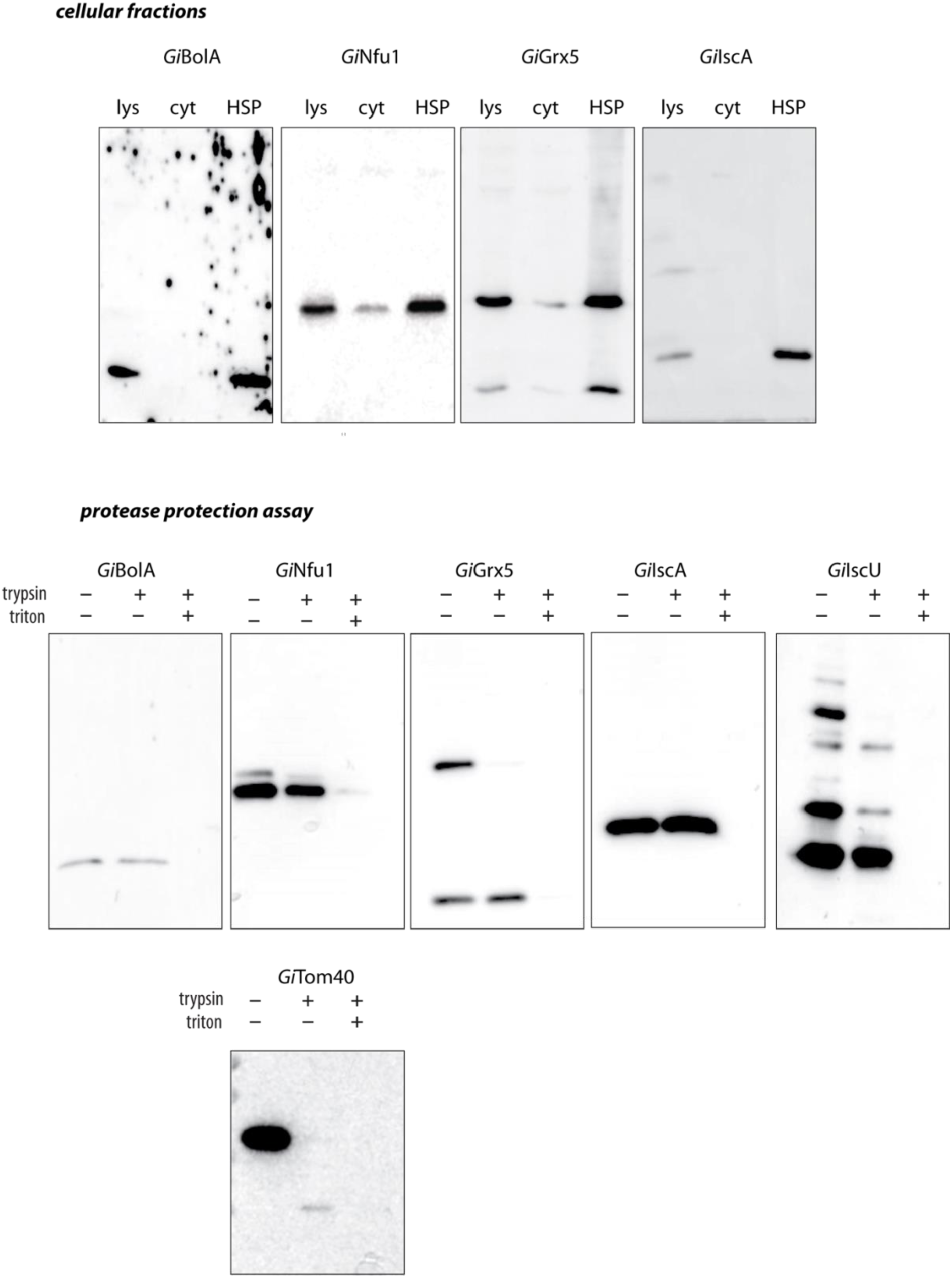
Full blots of cellular fractions and protease protection assay experiments.

**Supplementary Figure 3.**
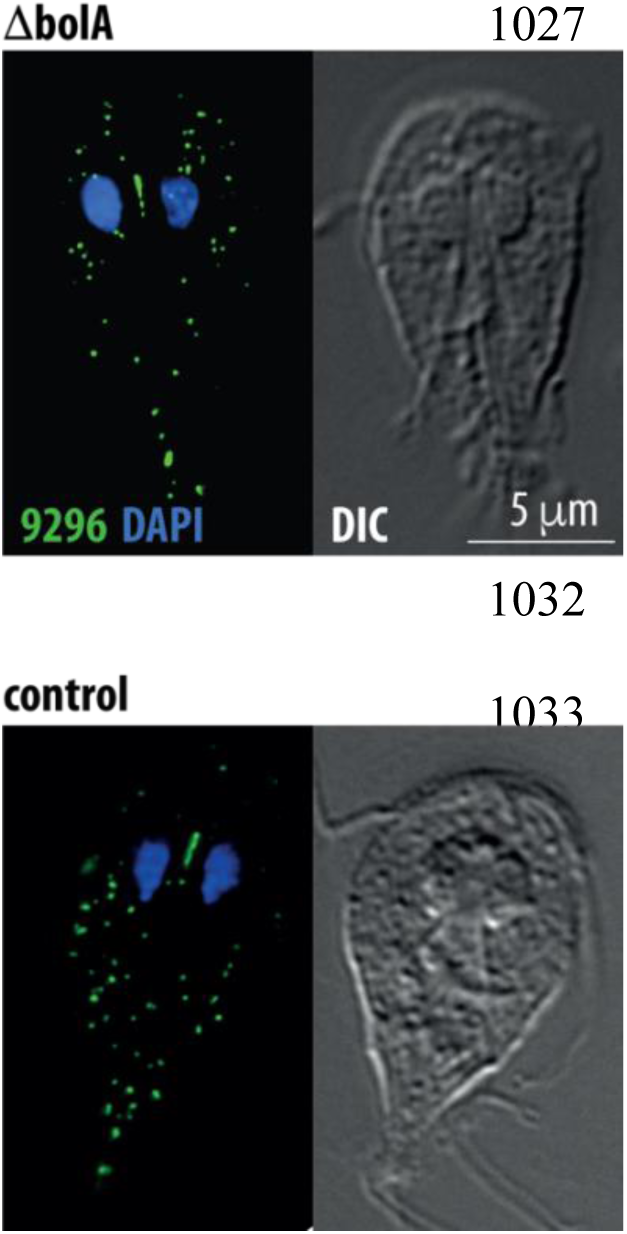
Mitosomal morphology and number is not affected by the removal of *bolA* gene. The exemplary image of mitosomes visualized by immunofluorescence microscopy in the ΔbolA and control (Cas9) cell lines. Mitosomes were detected by rabbit polyclonal antibody raised against GL50803_9296, the nuclei were stained with DAPI.

**Supplementary Table 1. Proteomic analysis of *Gi*BolA, *Gi*Grx5, *Gi*Nfu1 and *Gi*IscA pulldowns.** For all proteins, statistical analysis based upon the biological and technical triplicates are shown.

**Supplementary Table 2. ISC components of Metamonada**

**Supplementary Table 3. Fe-S proteins of *G. intestinalis*.**

**Supplementary Table 4. Proteomic analysis of ΔbolA cell line.**

**Supplementary Table 5. Primers used in the study.**

